# PCMD-1 bridges the centrioles and the PCM scaffold in *C. elegans*

**DOI:** 10.1101/2020.11.09.375865

**Authors:** Lisa Stenzel, Judith Mehler, Alina Schreiner, Sim Üstüner, Elisa Zuccoli, Esther Zanin, Tamara Mikeladze-Dvali

## Abstract

Correct cell division relies on the formation of a bipolar spindle. In animal cells, microtubule nucleation at the spindle poles is facilitated by the pericentriolar material (PCM), which assembles around a pair of centrioles. Although centrioles are essential for PCM assembly, proteins that anchor the PCM to the centrioles are less known. Here we investigate the molecular function of PCMD-1 in bridging the PCM and the centrioles in *Caenorhabditis elegans*.

We demonstrate that centrosomal recruitment of PCMD-1 is dependent on the outer centriolar protein SAS-7. While the most C-terminal part of PCMD-1 is sufficient to target it to the centrosome, the coiled-coil domain promotes its accumulation by facilitating self-interaction. We reveal that PCMD-1 is bridging the centrioles and PCM scaffold through protein-protein interactions with the PCM scaffold protein SPD-5, the mitotic kinase PLK-1 and the centriolar protein SAS-4. Using an ectopic translocation assay, we show that PCMD-1 is able to selectively recruit downstream PCM scaffold components to an ectopic location in the cell, indicating that PCMD-1 is sufficient to anchor the PCM scaffold proteins to the centrioles. Our work suggests that PCMD-1 is an essential functional bridge between the centrioles and the PCM.

## INTRODUCTION

Centrosomes are dynamic, non-membranous organelles that serve as the major microtubule-organizing centers in animal cells and are thus essential for biological processes ranging from polarity establishment to the orchestration of cell division. Centrosomes comprise a centriole pair and the surrounding pericentriolar material (PCM). The PCM dynamically changes in size and material properties during the cell cycle (Woodruff et al., 2015; 2017; Mittasch et al., 2020).

PCM expansion during mitosis facilitates bipolar spindle assembly. At the root of PCM expansion is a proteinaceous matrix that serves as a scaffold for the recruitment of regulatory proteins, including mitotic kinases and microtubule nucleators. In *C. elegans,* this scaffolding function is fulfilled by the self-assembly of the coiled-coil protein SPD-5 (functional homolog of Cdk5Rap2 in humans), which is controlled by phosphorylation through Polo-like kinase PLK-1 (homolog of PLK1 in humans) and the interaction with the coiled-coil protein SPD-2 (Cep192 homolog) (Hamill et al., 2002; Decker et al., 2011; Woodruff et al., 2015; 2017; Cabral et al., 2019). Our previous findings have revealed that PCMD-1, a protein with a predicted coiled-coil domain, regulates the spatial integrity of the PCM scaffold and together with SPD-2 is required for the recruitment of SPD-5 (Erpf et al., 2019). Centrioles serve as condensation centers for PCM proteins. During PCM expansion in mitosis, centrioles contribute to the growth and structural integrity of the PCM scaffold (Cabral et al., 2019). A limited set of centriolar core proteins have been described in *C. elegans* (O’Connell et al., 2001; Kirkham et al., 2003; Leidel and Gönczy, 2003; Dammermann et al., 2004; 2008; Delattre et al., 2004; 2006; Kemp et al., 2004; Pelletier et al., 2004; 2006; Leidel et al., 2005; Sugioka et al., 2017). From these proteins, SPD-2, SAS-4 (CPAP homolog) and SAS-7 have been proposed to functionally bridge the PCM and the centrioles (Varadarajan and Rusan, 2018). SAS-4, which localizes to the centrioles and the PCM, plays a critical role in microtubule assembly around the central tube of a forming centriole (Kirkham et al., 2003; Leidel and Gönczy, 2003; Dammermann et al., 2008; Delattre et al., 2006; Pelletier et al., 2006). SAS-7 facilitates the formation of paddlewheel structure on centriolar microtubules and recruits SPD-2, which in turn is needed for centriole duplication and mitotic PCM scaffold expansion (Sugioka et al., 2017).

We found that PCMD-1 is predominantly a centriolar protein (Erpf et al., 2019), yet its depletion affects SPD-5 recruitment and structural integrity of the PCM, raising the possibility that it functionally connects the PCM scaffold to the centrioles. However, the precise mechanisms of PCMD-1 centriolar targeting and the recruitment of other downstream proteins by PCMD-1 have still to be elucidated. Here we investigate the function of PCMD-1 in linking centrioles and PCM in *C. elegans* embryos. We found that centrosomal localization of PCMD-1 is dependent on the centriolar protein SAS-7. By analyzing different parts of PCMD-1, we found that the C-terminal part alone is sufficient for centrosomal targeting and that through PCMD-1 self-interaction, the coiled-coil domain promotes PCMD-1 accumulation at the centrosome. We demonstrate that PCMD-1 can physically interact with centriolar as well as PCM proteins, thus bridging the two components of the centrosome. Moreover, PCMD-1 is sufficient to selectively delocalize downstream PCM scaffold components to an ectopic location, supporting the notion that it can bridge centrioles and PCM.

## RESULTS

### SAS-7 recruits PCMD-1 to the centrioles in early embryos

Our previous analysis showed that most of the PCMD-1 protein localizes to the centrioles (Erpf et al., 2019). By recruiting SPD-5 to the centrosome, PCMD-1 functionally connects the PCM scaffold to the centrioles. This raises the question how PCMD-1 itself is anchored to the centrioles. PCMD-1 could be recruited to the centrosome via centriolar proteins. One candidate for such interaction is SAS-7 that forms the paddlewheels, the outermost centriolar structures known in *C. elegans* (Sugioka et al., 2017). We investigated the spatial relationship between PCMD-1 and SAS-7 by analyzing embryos expressing endogenously tagged GFP::PCMD-1 and RFP::SAS-7 using Lattice Structured Illumination Microscopy (SIM) (Figure 1A). We found that PCMD-1 and SAS-7 signals largely overlapped on both centrioles at spindle poles of mitotic blastomeres in early embryos (Figure 1A). We observed that one PCMD-1 focus had a higher accumulation of the SAS-7 signal, while the second one colocalized to a lesser extent with SAS-7, consistent with its function in centriole maturation and daughter centriole formation (arrowheads, Figure 1A) (Sugioka et al., 2017). This observation prompted us to test whether SAS-7 localization is dependent on PCMD-1. Live-cell imaging of GFP::SAS-7 in the *pcmd-1(t3421)* mutant animals revealed that GFP::SAS-7 was always detected at centrioles, similar to control embryos (Figure 1B). We concluded that PCMD-1 is not involved in the recruitment of SAS-7 to the centrioles. To test inversely whether SAS-7 is required for PCMD-1 recruitment to the centrosome, we expressed *gfp::pcmd-1* in *sas-7(or452)* mutant animals. While GFP::PCMD-1 at the centrosome was apparent in all control embryos, centrosomal GFP::PCMD-1 signal was highly reduced, almost undetectable in *gfp::pcmd-1*;*sas-7(or452)* one-cell embryos (Figure S1A). Interestingly, a weak GFP::PCMD-1 signal was consistently detected at sperm-derived centrioles during pronuclear migration (Figures S1A). Shortly thereafter, this signal fell below the detection limit in 5 out of 16 centrosomes (Figures S1B). In the remaining embryos, some negligible GFP signal remained at the centrosome, reflecting the hypomorphic nature of the *sas-7(or452)* allele (data not shown) (Sugioka et al., 2017). As an alternative means to address the SAS-7 dependent localization of PCMD-1, we performed immunostaining using antibodies against GFP and SAS-4 to mark the centrioles. In one-cell control embryos, all centrosomal SAS-4 foci colocalized with a clear GFP::PCMD-1 signal (Figure 1C). In contrast, in none of the *gfp::pcmd-1*;*sas-7(or452)* mutant one-cell embryos, did centrosomal SAS-4 foci colocalized with a GFP::PCMD-1 signal (Figure 1C). Note that centrosomal GFP::PCMD-1 signal was detectable in later multicellular embryos (Figure S1C). Thus, SAS-7 is necessary to recruit and maintain PCMD-1 during the first cell divisions.

**Figure 1.**
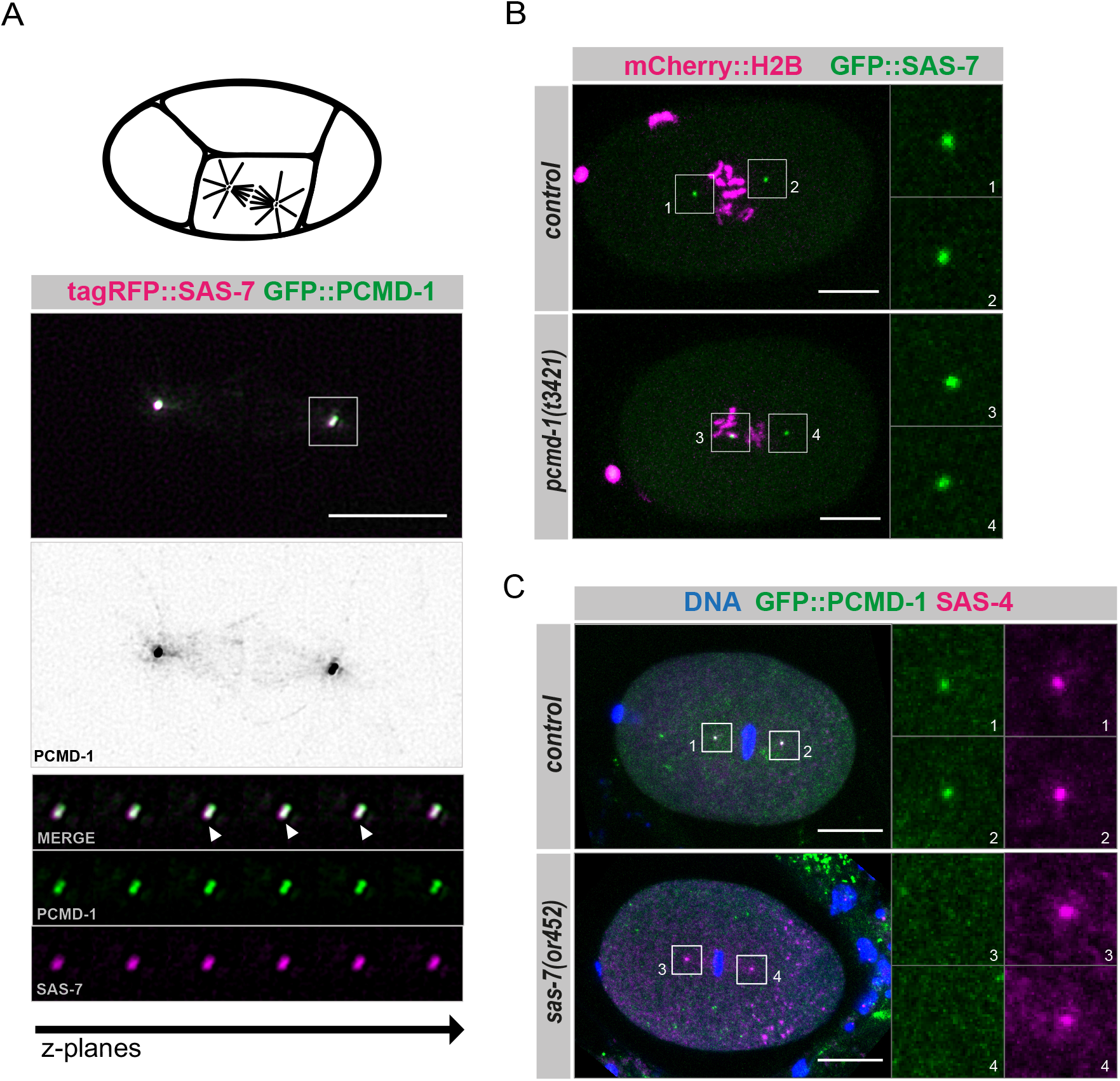
SAS-7 recruits PCMD-1 to the centrioles. (A) Mitotic centrosomes in a 4-cell embryo expressing GFP::PCMD-1 and tagRFP::SAS-7 (n=6 centrosomes). Note that some GFP signal decorated astral and kinetochore microtubules of the mitotic spindle (middle panel). Lower panels represent a montage of different Z-planes spanning a centriole pair. Arrowheads indicate Z-planes where a higher accumulation of the SAS-7 signal is visible at one of the two centrioles. (B) Stills of time-lapse spinning disc confocal images of *mCherry::h2b;gfp::sas-7* (n=16) and *pcmd-1(t3421);mCherry::h2b;gfp::sas-7* (n=12) embryos during nuclear envelope break-down (NEB). Insets represent centrosomes. (C) Representative confocal images of fixed *gfp::pcmd-1* (n=16) and *gfp::pcmd-1;sas-7(or452)* (n=6) one-cell embryos in prometaphase stained for DNA, GFP and SAS-4. Insets represent single channels of the centrioles. In all panels, scale bars are 10 μm.

### PCMD-1 bridges centriolar and PCM scaffold proteins

The genetic dependency of PCMD-1 centrosomal localization on SAS-7 raises the possibility that PCMD-1 is recruited to the centriole by direct interaction with SAS-7. To address if PCMD-1 and SAS-7 interact with each other and other centrosomal proteins such as SAS-4, SPD-2, SPD-5, and PLK-1, we performed a candidate-based yeast two-hybrid screen with a dual reporter system based on LexA expression (Fields and Song, 1989). The readouts of positive interactions were both growth on leucine-lacking medium and expression of a GFP-reporter. We generated bait-plasmids containing full-length cDNAs of SAS-7 and PCMD-1 and prey-plasmids for the following candidate proteins: SAS-4, SAS-7, PCMD-1, SPD-2, SPD-5, and PLK-1. After expressing the bait- and prey-plasmids in yeast, growth and expression of the GFP-reporter were monitored on day 3 and 5. The interacting partners p53 and LTA, a positive control provided by the manufacturer, showed growth and GFP expression on day 3. Hereafter, we categorize combinations of proteins showing both readouts on day 3 as strong interactors and one or both readouts on day 5 as weak interactors.

Previous yeast two-hybrid screens identified SPD-2 as a SAS-7 binding protein (Li et al., 2004; Boxem et al., 2008; Sugioka et al., 2017). We used this interaction to validate our yeast two-hybrid assay for centrosomal proteins. As previously reported, we found that SAS-7 strongly interacts with SPD-2 and we could also recapitulate a weaker interaction of SAS-7 with SAS-4 (Figures 2A, 2B) (Boxem et al., 2008). However, we could not detect any interaction between the SAS-7 bait and PCMD-1 or SAS-7 prey. Since we did not detect an interaction between PCMD-1 and SAS-7, we tested whether PCMD-1 as bait could interact with SAS-7, SAS-4, or SPD-2. No growth was observed for colonies expressing PCMD-1 and SAS-7 or PCMD-1 and SPD-2, but we could detect strong protein-protein interaction of PCMD-1 with SAS-4. Next, we tested whether PCMD-1 could bind to the PCM scaffold protein SPD-5, the kinase PLK-1 or itself. We could detect strong protein-protein interaction of PCMD-1 with itself and weaker interaction with both SPD-5 and PLK-1. In turn, SAS-7 as a prey did not interact with SPD-5 but showed a very weak growth of colonies when expressed together with PLK-1 (Figures 2A, 2B), although the expression of the GFP reporter was below detection level.

**Figure 2.**
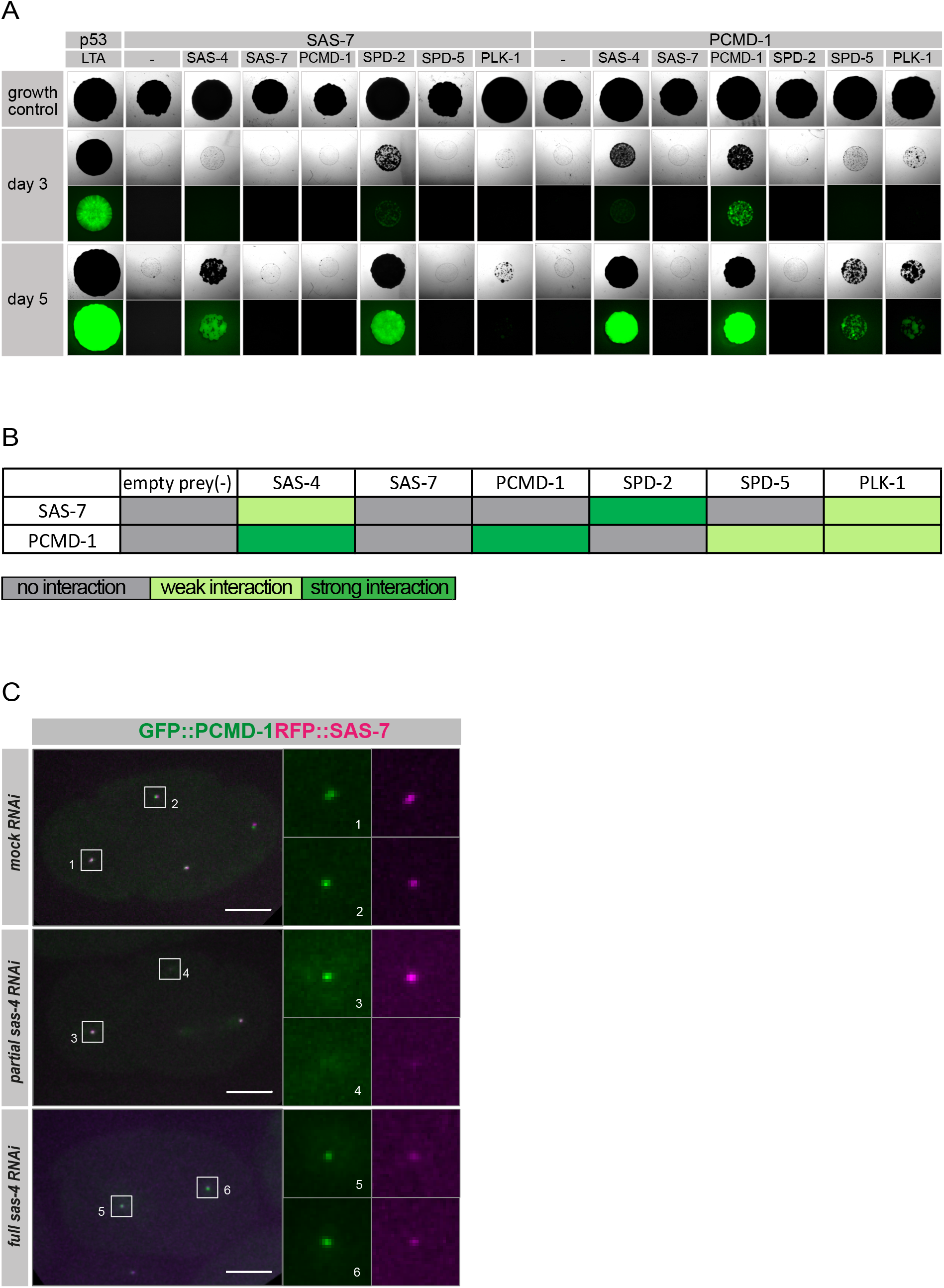
PCMD-1 interacts with centriolar and PCM scaffold proteins. (A) Images of representative yeast two-hybrid colonies. Interaction of bait proteins SAS-7 and PCMD-1 with centrosomal proteins SAS-7, SAS-4, PCMD-1, SPD-2, SPD-5, and PLK-1 as preys, respectively. The top panel represents growth control on medium with leucine, middle panels represent colonies screened on day 3 for growth on leucine-lacking medium and expression of the GFP-reporter, and bottom panels represent colonies screened on day 5 for growth on leucine-lacking medium and expression of the GFP-reporter. (B) Summary of protein-protein interactions categorized by the strength of their interactions. (C) Stills of time-lapse spinning disc confocal images of dividing GFP::PCMD-1; tagRFP::SAS-7 expressing 2-cell embryos treated with *mock* or *sas-4 RNAi*. n=8, n=16 and n=11 for *mock,* partial and full *sas-4* RNAi-treated embryos, respectively. Insets represent single channels of the centrioles. In all panels, scale bars are 10 μm.

Taken together, our findings in the yeast two-hybrid assay do not support the hypothesis that SAS-7 directly interacts with PCMD-1 or that the interaction occurs through SPD-2. Interestingly, both proteins seem to bind to SAS-4 (Boxem et al., 2008). This could in principle suggest that SAS-4 is mediating the dependency of PCMD-1 on SAS-7.

To test this hypothesis, we treated embryos with *sas-4*(*RNAi)* and monitored GFP::PCMD-1 and tagRFP::SAS-7 recruitment to the site of a prospective daughter centriole in 2-cell embryos. Centriole duplication fails SAS-4 depleted embryos. As a consequence, monopolar spindles form in the blastomeres of a 2-cell embryo (Kirkham et al., 2003; Leidel and Gönczy, 2003). In strong *sas-4*(*RNAi)* embryos PCMD-1 and SAS-7 were only detected at the mother centrioles (Figure 2C). In partial *sas-4*(*RNAi)* embryos, where in some cases structurally incomplete daughter centrioles form (Kirkham et al., 2003), we either found a weak signal of both or none of the two proteins. This is in contrast to *mock(RNAi)*-treated embryos, where a strong signal of GFP::PCMD-1 and tagRFP::SAS-7 is detected at both spindle poles (Figure 2C). Our results suggest, that both PCMD-1 and SAS-7 need some functional SAS-4 to be recruited to the nascent daughter centrioles. However, we and others have shown, that in *sas-7(or452)* mutant embryos, where no PCMD-1 is found at the centrosome, SAS-4 foci are still present (Figure 1C) (Sugioka et al., 2017). Therefore, it is highly unlikely that SAS-4 is accountable for the loss of PCMD-1 centrosomal localization in *sas-7(or452)* embryos.

In summary, PCMD-1 binds to the centriolar protein SAS-4, the PCM protein SPD-5, the mitotic kinase PLK-1, and to itself. Therefore, we propose that PCMD-1 physically bridges the centrioles and the PCM scaffold.

### PCMD-1 is sufficient to recruit SPD-5 and PLK-1 to an ectopic location

We previously demonstrated that PCMD-1 is required to recruit SPD-5 to the centrosome (Erpf et al., 2019). This finding is strongly supported by our protein-protein interaction data, which indicates that PCMD-1 bridges centriolar and PCM proteins. Therefore, we asked whether PCMD-1 is also sufficient to recruit SPD-5. To address this question, we set up a ‘translocation assay’ by targeting PCMD-1 to an ectopic location in the cell and testing whether PCMD-1 is capable to recruit SPD-5 to this cellular location. To tether PCMD-1 to the plasma membrane, we fused the mKate2::PCMD-1 reporter to the plcδ1PH*-*domain and expressed it under a heat-shock promoter. Upon heat-shock PH::mKate2::PCMD-1 was expressed and reliably translocated to the plasma membrane (Figures 3A, 3B).

**Figure 3.**
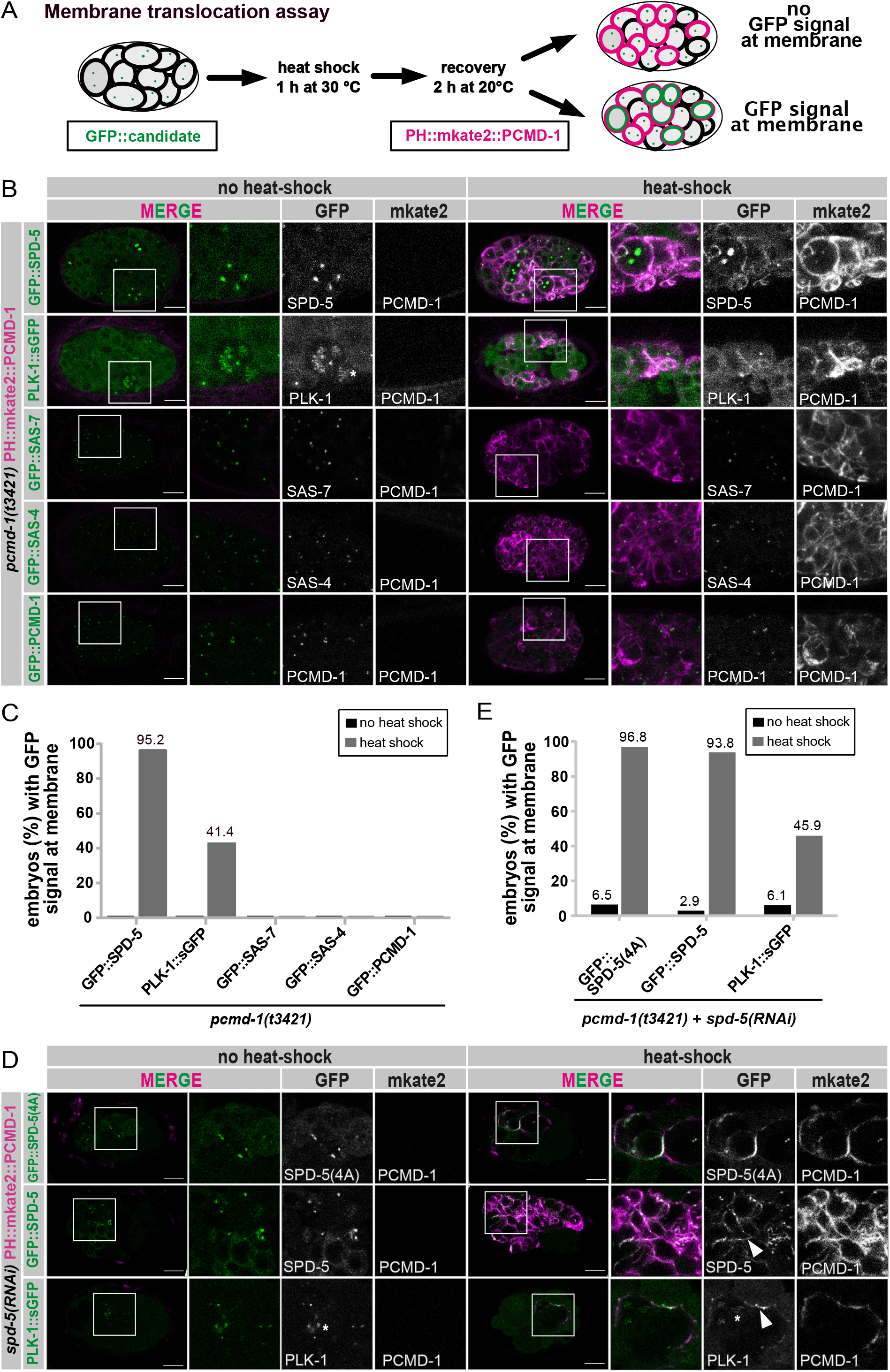
PCMD-1 targets SPD-5 and PLK-1 to the plasma membrane. (A) Schematic representation of the ‘translocation assay’. After 1h heat shock to induce expression and 2h recovery, PH::mKate2::PCMD-1 is expressed and binds to the plasma membrane of a multicellular embryo. If PH::mKate2::PCMD-1 recruits GFP-tagged candidate proteins, they will also localize to the plasma membrane (bottom embryo). If PH::mKate2::PCMD-1 is not able to recruit, the localization of the GFP-tagged candidate proteins will not change upon PH::mKate2::PCMD-1 expression (top embryo). (B) Representative multicellular embryos of the ‘translocations assay’ for GFP::SPD-5 (n=17 no heat shock; n=21 heat shock), PLK-1::sGFP (n=21 no heat shock; n=29 heat shock), GFP::SAS-7 (n=20 no heat shock; n=23 heat shock), GFP::SAS-4 (n=28 no heat shock; n=28 heat shock) and GFP::PCMD-1 (n=25 no heat shock; n=27 heat shock) fusion proteins in the *pcmd-1(t3421)* background with and without heat-shock. Selected regions are enlarged and shown as merge and single channels. Note that PLK-1::sGFP signal at the plasma membrane is less intense than GFP::SPD-5. Scale bars are 10 μm. (C) Quantification of (B); percentage of embryos (%) with GFP signal at the membrane after heat shock in the *pcmd-1(t3421)* background (n= total number of embryos analyzed). (D) Representative multicellular embryos of the ‘translocations assay’ using GFP::SPD-5 (n=31 no heat shock; n=31 heat shock), GFP::SPD-5(4A) (n=34 no heat shock; n=32 heat shock), and PLK-1::sGFP (n=33 no heat shock; n=37 heat shock) in a *pcmd-1(t3421)* background and treated by SPD-5 RNAi. Selected regions are enlarged and shown as merge and single channels. Arrowheads indicate the membrane-localized GFP signal. Asterisk indicated kinetochore localization of PLK-1::sGFP. Scale bars are 10 μm. (E) Quantification of (D); percentage of embryos (%) with GFP signal at the membrane after heat shock (n= total number of embryos analyzed).

We tested whether PH::mKate2::PCMD-1 was sufficient to recruit GFP::SPD-5 in the *pcmd-1(t3421)* mutants and in the presence of wild-type PCMD-1. In *pcmd-1(t3421)* animals, recruitment of SPD-5 to the centrosome is compromised due to the lack of endogenous PCMD-1, therefore more GFP::SPD-5 is expected in the cytoplasm (Erpf et al., 2019). In control embryos without heat shock GFP::SPD-5 was never detected at the plasma membrane. However, after heat shock in 95.2% of *pcmd-1(t3421)* and 68.4% of wild-type embryos PH::mKate2::PCMD-1 recruited GFP::SPD-5 to the plasma membrane (Figures 3B, 3C and Figures S2A, S2B).

Since PLK-1 phosphorylates SPD-5 and also interacts with PCMD-1 in the yeast two-hybrid assay, we tested if PH::mKate2::PCMD-1 was sufficient to recruit PLK-1::sGFP to the plasma membrane. Similar to GFP::SPD-5, in 41.4% of *pcmd-1(t3421)* and 26.3% of wild-type embryos PLK-1::sGFP was recruited to the plasma membrane (Figures 3B, 3C and Figures S2A, S2B). Note that the PLK-1::sGFP signal at the membrane was much weaker compared to the GFP::SPD-5 signal.

Since PCMD-1 is sufficient to recruit SPD-5 as well as PLK-1 to an ectopic location, we wondered whether the membrane-bound pool of SPD-5 required the phosphorylation by PLK-1, which has been shown to be essential for PCM maturation (Woodruff et al., 2015). For this we used a strain carrying GFP::SPD-5^4A^, in which the four PLK-1 phosphorylation sites needed for the maturation and expansion of the mitotic scaffold were substituted by alanines (Woodruff et al., 2015). We found that PH::mKate2::PCMD-1 was still able to recruit GFP::SPD-5^4A^ to the membrane (93.8 %, Figures 3D, 3E), indicating that the membrane-bound GFP::SPD-5 pool represents the PCM core fraction of SPD-5, rather than the expandable mitotic scaffold. Reciprocally, PLK-1::sGFP recruitment still took place in a SPD-5 RNAi background (45.9% Figures 3D, 3E). Thus, PCMD-1 can independently recruit PLK-1 and SPD-5.

Interestingly, PH::mKate2::PCMD-1 was unable to recruit the centriolar proteins GFP::SAS-4, GFP::SAS-7 or GFP::PCMD-1 in any of the analyzed *pcmd-1(t3421)* embryos (Figures 3B, 3C). This was surprising, especially for SAS-4 and PCMD-1, since their interaction in the yeast two-hybrid assay was very strong. Therefore, the ability of PCMD-1 to ectopically anchor proteins to the plasma membrane is specific to PCM proteins.

These findings suggest that PCMD-1 would also be sufficient to recruit the PCM scaffold to the centrosome in the *C. elegans* one-cell embryo. Since PCMD-1 is sufficient to recruit SPD-5, we next asked whether PCMD-1 is loaded onto centrosomes before SPD-5. In the *C. elegans* zygote, SPD-5 is recruited to the sperm-derived centrioles after the completion of meiosis II of the female pronucleus and concomitant with the ability of the centrosome to nucleate microtubules (McNally et al., 2012). This paradigm allows us to investigate whether PCMD-1 is loaded to the centrioles prior to SPD-5 recruitment using marked mating experiments where only the sperm or the oocyte express a fluorescent marker. First, we tested whether paternal GFP::PCMD-1 could be detected at the centrosome after fertilization. For this we mated *fog-2(q71)* females lacking sperms with GFP::PCMD-1 expressing males, thus labelling sperm centrioles (Figure S3A) (Erpf et al., 2019). We analyzed the embryos during the first mitotic division. We could not detect GFP::PCMD-1 at centrioles, in any of the analyzed embryos. Second, we mated GFP::PCMD-1 females, treated with *fem-1* RNAi to block sperm production, with control *fog-2(n71)* males with unlabeled sperm centrioles. We found that GFP::PCMD-1 signal was detected at the centrosomes in all analyzed embryos during the first mitotic division (Figure S3B). These results suggest a turnover of sperm derived PCMD-1 after fertilization.

To determine when exactly maternal PCMD-1 is recruited to the centriole after fertilization and to temporally map its loading with respect to SPD-5, we immuno-stained embryos from GFP::PCMD-1 females mated with males with unlabeled sperm centrioles (Figures S3B, S3C), with antibodies against SPD-5 and against GFP. In meiosis I embryos, neither GFP::PCMD-1 nor SPD-5 foci were present at the centrioles (Figure S3D), indicating that maternal GFP::PCMD-1 was not yet incorporated in the centrioles. During meiosis II we found that a GFP::PCMD-1 focus, without any detectable SPD-5, was visible at 90.5% of sperm centrioles (Figures S3C, S3D). In the remaining 9.5% of embryos, categorized as early meiosis II, neither GFP::PCMD-1 nor SPD-5 foci were present (Figure S3C). Therefore, we conclude that maternal GFP::PCMD-1 is recruited to the sperm centrioles at meiosis II. After meiosis II, when sperm pronuclei are decondensed, SPD-5 colocalized with GFP::PCMD-1 at centrosomes in 87% of embryos (Figures S3C, S3D). We never observed embryos with centrosomes only labeled by SPD-5.

In summary, our results are consistent with a model in which maternal GFP::PCMD-1 is recruited to the sperm derived centrioles shortly after fertilization and prior to SPD-5 loading. We suggest that PCMD-1 is sufficient to anchor SPD-5 to the centrosome and to recruit PLK-1 to initiate centrosome maturation. These findings strengthen the hypothesis that PCMD-1 is bridging centriolar and PCM proteins.

### The coiled-coil domain of PCMD-1 promotes its loading to the centrosome and PCM scaffold formation

To determine how PCMD-1 is anchored to the centrosome, we next examined which part of the protein is necessary for its centrosomal targeting. PCMD-1 is predicted to have a single coiled-coil domain and multiple Intrinsically Disordered Regions (IDRs), which partially overlap with low complexity regions (Figure 4A) (UniProt Consortium, 2019; Schultz et al., 2000; Letunic and Bork, 2018). Coiled-coil domains often mediate protein-protein interactions, including oligomerization and these interactions can have regulatory functions for centrosomal proteins (Leidel et al., 2005; Kitagawa et al., 2011; Qiao et al., 2012; Hilbert et al., 2013; Lettman et al., 2013; Rogala et al., 2015). Therefore, we set out to investigate the function of the coiled-coil domain in PCMD-1.

**Figure 4.**
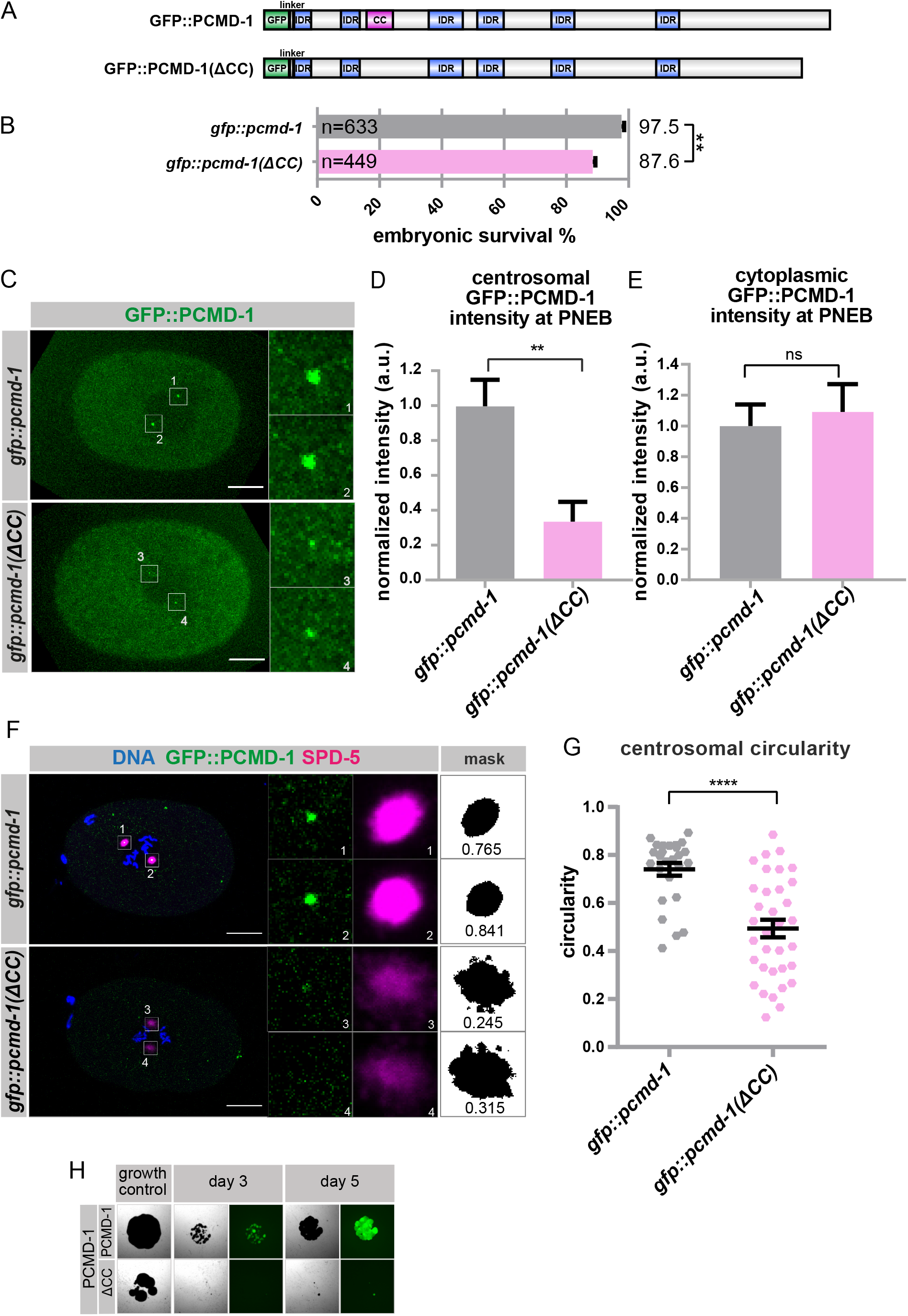
The coiled-coil domain promotes PCMD-1 accumulation at the centrosome and PCM scaffold integrity. (A) Schematic representation of the domain structure of endogenously tagged GFP::PCMD-1 protein (aa 2-630), with predictions of the coiled-coil domain (CC) and Intrinsically Disordered Regions (IDRs) (top). Schematic representation of the domain structure of a truncated version, with the deleted coiled-coil domain (Δ86-117). All domains except GFP are represented to the correct ratio. (B) Embryonic survival (%) of *gfp::pcmd-1* and *gfp::pcmd-1(*Δ*CC)* expressing animals at 25°C. Kruskal-Wallis test. n=number of analyzed embryos. (C) Stills of time-lapse imaging of embryos expressing *gfp::pcmd-1* (n=7) and *gfp::pcmd-1(*Δ*CC)* (n=9) at NEB. Centrosomal areas were determined by DIC imaging and are shown enlarged for the GFP::PCMD-1 signal. Also, see movies 1 and 2. (D) Normalized centrosomal GFP signal intensities in embryos expressing *gfp::pcmd-1* (n=14, 1.00±0.15 arbitrary units (a.u.), mean±s.e.m.) and *gfp::pcmd-1(*Δ*CC)* (n=18, 0.34±0.11 a.u., mean±s.e.m.) at NEB. Wilcoxon test. n=number of analyzed centrosomes. (E) Normalized cytoplasmic GFP signal intensities in embryos expressing *gfp::pcmd-1* (n=7, 1±0.14 a.u., mean±s.e.m.) and *gfp::pcmd-1(*Δ*CC)* (n=9, 1.09±0.18 a.u., mean±s.e.m.) at NEB. Two-sample t-test. n=number of analyzed embryos. (F) Representative images of fixed embryos of the indicated genotype stained for DNA, GFP and SPD-5. Enlarged are centrosomes in individual channels, corresponding mask and values of centrosomal circularity. (G) Quantification of SPD-5 circularity using the indicated formula in *gfp::pcmd-1* (n=26, 0.74±0.03, mean±s.e.m.) and *gfp::pcmd-1(*Δ*CC)* (n=34, 0.49±0.04, mean±s.e.m.) embryos. Images of SPD-5 areas of centrosome 1 and 3. Kruskal-Wallis test, n=number of analyzed centrosomes. (H) Images of representative yeast two-hybrid colonies. Interaction of bait proteins PCMD-1 with PCMD-1 and PCMD-1(ΔCC) as preys. In all panels error bars denote s.e.m.. P-values represent **p<0.01, ****p<0.0001, ns = not siginificant. Scale bars are 10 μm.

To examine the function of the coiled-coil domain, we deleted the sequence from E86 including F118, predicted as the coiled-coil domain by the COILS program (Lupas et al., 1991; Lupas, 1996), using CRISPR/Cas9 in the context of the in locus-tagged GFP::PCMD-1 protein. We refer to this deletion as *gfp::pcmd-1*(*ΔCC*) (Figure 4A). In a lethality test 97.5% of the *gfp::pcmd-1* embryos and 87.6% of the *gfp::pcmd-1(ΔCC)* embryos survived at 25°C (Figure 4B, Table 1). Thus, the deletion of the predicted coiled-coil domain slightly compromised embryonic viability.

**Table 1.** Embryonic viability counts at different temperatures

| Genotype | Temp. | Embryonic viability<br>(%, mean ± s.e.m.) | worms | eggs |
| --- | --- | --- | --- | --- |
| *pcmd-1(syb486[gfp::pcmd-1])**l* | 25 °C | 97.45 ± 0.06 | 34 | 633 |
| *pcmd-1(syb1285 syb486[gfp::pcmd-1(Δcc)])**l* | 25 °C | 87.63 ± 0.03 | 50 | 449 |
| *pcmd-1(t3421); mikSi6[Pmai-2:GFP::PCMD-1]* *ll* | 25 °C | 96.32 ± 0.47 | 111 | 3399 |
| *pcmd-1(t3421)* *l* | 25 °C | 0.00 ± 0.00 | 94 | 3163 |
| *pcmd-1(t3421); mikSi14[Pmai-2:GFP::PCMD-1_Nter(1-117)]* *ll* | 25 °C | 0.33 ± 0.19 | 48 | 1718 |
| *pcmd-1(t3421); mikSi15[Pmai-2:GFP::PCMD-1_Cter(118-630)]* *ll* | 25 °C | 90.8 ± 1.09 | 50 | 2159 |
| *pcmd-1(t3421); mikSi16[Pmai-2:GFP::PCMD-1_C1(118-342)]* *ll* | 25 °C | 0.00 ± 0.00 | 32 | 1437 |
| *pcmd-1(t3421); mikSi17[Pmai-2:GFP::PCMD-1_C2(342-630)]* *ll* | 25 °C | 0.00 ± 0.00 | 29 | 1157 |
| *pcmd-1(t3421); mikSi6[Pmai-2:GFP::PCMD-1]* *ll* | 15 °C | 93.62 ± 1.27 | 46 | 1125 |
| *pcmd-1(t3421)* *l* | 15 °C | 40.99 ± 2.68 | 46 | 2063 |
| *pcmd-1(t3421); mikSi14[Pmai-2:GFP::PCMD-1_Nter(1-117)]* *ll* | 15 °C | 28.03 ± 1.72 | 64 | 1086 |

Next, we investigated if the GFP::PCMD-1(ΔCC) protein could localize to the centrosome. We performed live-cell imaging on worms expressing GFP::PCMD-1 and GFP::PCMD-1(ΔCC). While GFP::PCMD-1 efficiently localized to centrosomes in all analyzed embryos, the GFP::PCMD-1(ΔCC) signal on average appeared much weaker, even falling below the detection limit at 3 out of 18 centrosomes (Figure 4C). Measuring the mean centrosomal GFP signal intensity confirmed that PCMD-1 without the coiled-coil domain was significantly reduced at centrosomes in comparison to wild-type PCMD-1, while cytoplasmic levels remained similar to the wild type (Figures 4D, 4E). Thus, the coiled-coil domain is necessary for efficient centrosomal loading of PCMD-1, but is not essential for the viability of the embryos. This raises the question how embryos with reduced centrosomal PCMD-1 levels can divide. To investigate whether these animals could still recruit the PCM scaffold, we immuno-stained GFP::PCMD-1(ΔCC) embryos using antibodies against GFP and SPD-5. We found that compared to controls, the GFP::PCMD-1(ΔCC) signal was largely reduced but still detectable in most embryos. SPD-5 was still recruited to the centrosome in all analyzed embryos, even in embryos where GFP::PCMD-1(ΔCC) was almost undetectable (Figure 4F). Intriguingly, in the absence of the coiled-coil domain, the SPD-5 centrosome matrix appeared to be much more dispersed and disorganized (Figure 4F). Mean centrosome circularity values significantly drop in *gfp::pcmd-1(*Δ*CC)* animals (Figure 4G).

Since coiled-coil domains are often implemented in the oligomerization of centrosomal proteins, we asked the question whether PCMD-1 self-interaction was compromised in the absence of the coiled-coil domain. To this end, we expressed PCMD-1(ΔCC) as a prey plasmid in combination with the PCMD-1 full-length bait plasmid in the yeast two-hybrid system. PCMD-1 self-interaction was abolished in the absence of the coiled-coil domain (Figure 4H). We propose that the coiled-coil domain facilitates PCMD-1 oligomerization and thereby promotes efficient PCMD-1 accumulation at the centrosome and the maintenance of PCM scaffold integrity.

### The C-terminal region of PCMD-1 is sufficient to target the protein to the centrosome

The fact that in some of the embryos GFP::PCMD-1(ΔCC) could still be recruited to the centrosome indicates that protein domains other than the predicted coiled-coil domain might play a role in centrosomal anchoring. To map which part of the protein is involved, we used a previously established single copy replacement system (Erpf et al., 2019). *pcmd-1(t3421)* mutant animals are 100% lethal at 25°C (Figures 5A, 5B, Table 1). Reconstituting a single copy of the PCMD-1 cDNA under the regulatory elements of the *mai-2* gene restores survival rates to 95.6% (Figure 5B, Table 1). We used this assay to test the functionality of different truncations of the PCMD-1 protein. GFP::PCMD-1(N) spans the region E002-N117, including the first two IDRs and the coiled-coil domain. GFP::PCMD-1(C) comprises amino acids F118 to the stop codon, spanning the remaining IDRs (Figure 5A, Table 1). In the survival assay GFP::PCMD-1(C) could restore embryonic lethality of *pcmd-1(t3421)* to 90.8% survival, while GFP::PCMD-1(N) was not sufficient to rescue viability (0.3%) (Figure 5B, Table 1). Interestingly, GFP::PCMD-1(N) had even a negative effect on the viability of *pcmd-1(t3421)* at the permissive temperature of 15°C, reducing it from 41% to 28% (Figure S4A, Table 1).

**Figure 5.**
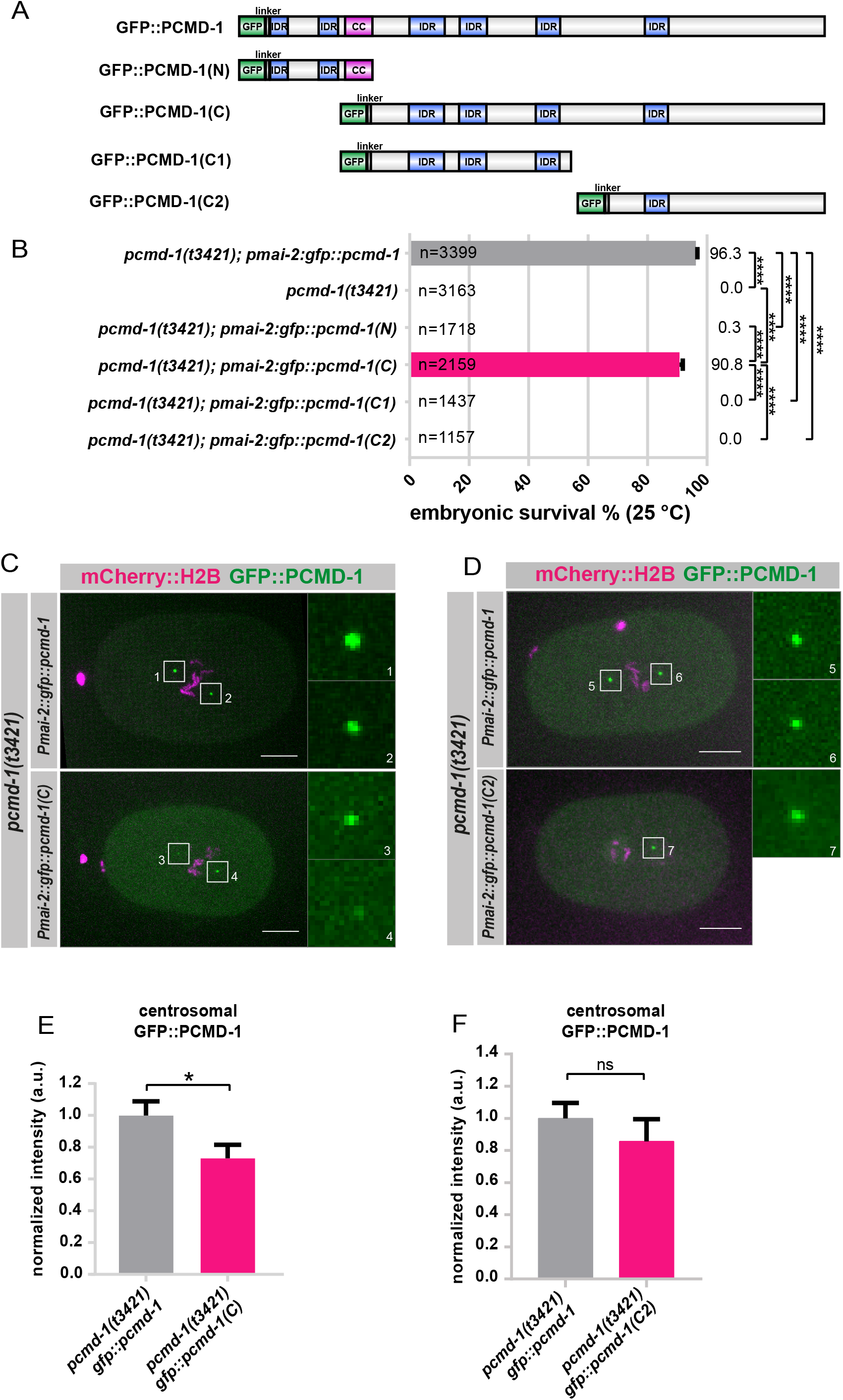
The most C-terminal region of PCMD-1 is sufficient to target the protein to the centrosome. (A) Domain structure of different GFP tagged PCMD-1 constructs. Top shows the GFP::PCMD-1 full-length protein, middle the GFP::PCMD-1(N) which spans the N-terminus (aa 1-117) including the predicted coiled-coil domain (CC) and bottom GFP::PCMD-1(C) represents the C-terminus (aa 118-630), excluding the coiled-coil domain. GFP::PCMD-1(C1) spans (aa 118-342) and GFP::PCMD-1(C2) spans (aa 343-630). All domains except GFP are represented to the correct ratio. (B) Embryonic survival (%) of *Pmai-2::gfp::pcmd-1, Pmai-2::gfp::pcmd-1(N), Pmai-2::gfp::pcmd-1(C), Pmai-2::gfp::pcmd-1(C1)* and *Pmai-2::gfp::pcmd-1(C2)* in the *pcmd-1(t3421)* background at 25°C. Multiple Comparison with Kruskal Wallis test and post-hoc Dunn’s test adjusted with Holm correction. Results with no significance are not indicated in the graph, n=number of analyzed embryos. (C) Stills of time-lapse imaging of embryos expressing *pcmd-1(t3421);Pmai-2::gfp::pcmd-1; Ppie-1::mCherry::h2b* (n=12) and *pcmd-1(t3421);Pmai-2::gfp::pcmd-1(C);Ppie-1::mCherry::h2b* (n=10) at NEB. Enlarged are the two centrosomes. (D) Stills of time-lapse imaging of embryos expressing *pcmd-1(t3421);Pmai-2::gfp::pcmd-1; Ppie-1::mCherry::h2b* (n=9) and *pcmd-1(t3421);Pmai-2::gfp::pcmd-1(C2),Ppie-1::mCherry::h2b* (n=9) at NEB. Enlarged are the two centrosomes. (E) Normalized centrosomal GFP signal intensities of embryos expressing *pcmd-1(t3421);Pmai-2::gfp::pcmd-1; Ppie-1::mCherry::h2b* (n=24, 1.00±0.09 a.u., mean±s.e.m.) and *pcmd-1(t3421);Pmai-2::gfp::pcmd-1(C);Ppie-1::mCherry::h2b* (n=20, 0.73±0.09 a.u., mean± s.e.m.) at NEB. Wilcoxontest. n=number of analyzed centrosomes. (F) Normalized centrosomal GFP signal intensities of embryos expressing *pcmd-1(t3421);Pmai-2::gfp::pcmd-1; Ppie-1::mCherry::h2b* (n=18, 1±0.1 a.u., mean±s.e.m.) and *pcmd-1(t3421);Pmai-2::gfp::pcmd-1(C2);Ppie-1::mCherry::h2b* (n=15, 0.86±0.14 a.u., mean±s.e.m.) at NEB. Two Sample t-test. n=number of analyzed centrosomes. In all panels error bars denote s.e.m.. P-values represent * p<0.05, **** p<0.0001, ns= not significant. Scale bars are 10 μm.

Next, we assessed the ability of these constructs to localize to the centrosome by live-cell imaging. Centrosomal GFP signal was detected in all GFP::PCMD-1(C) embryos, albeit the GFP signal intensities were significantly reduced compared to control animals (Figures 5C, 5E). In contrast, we could not detect any GFP signal at the centrosome in embryos expressing the GFP::PCMD-1(N) constructs, even though the cytoplasmic levels were significantly higher (Figures S4B, S4C). Therefore, the C-terminal part of PCMD-1, excluding the coiled-coil domain and the first two predicted IDRs is sufficient for PCMD-1 anchoring to the centrosome.

To further map the part of PCMD-1 that targets the protein to the centrosome, we subdivided the C-terminal part into two fragments spanning F118-D342 (C1) and G343-stop codon (C2) (Figure 5A). In the survival assay, neither GFP::PCMD-1(C1) nor GFP::PCMD-1(C2) could rescue embryonic lethality of *pcmd-1(t3421)* (Figure 5B, Table 1). However, in contrast to GFP::PCMD-1(C1), which did not exhibit any cellular localization pattern (Figure S4C), GFP::PCMD-1(C2) localized to the centrosome in all analyzed embryos (Figures 5D, 5F). Similar to the centrosomes in the embryo, only GFP::PCMD-1, GFP::PCMD-1(C) and GFP::PCMD-1(C2) localized to the ciliary base of adult animals (Figure S4D). Therefore, the C-terminal part of PCMD-1, including the last IDR, is sufficient to target PCMD-1 to the centrosome and the ciliary base.

## DISCUSSION

In this study, we reveal the mechanism by which PCMD-1 anchors the PCM to the centriole. We demonstrate that PCMD-1 directly interacts with the downstream PCM scaffold protein SPD-5 and the kinase PKL-1. Furthermore, PCMD-1 is sufficient to recruit SPD-5 and PKL-1 to an ectopic cellular location. Intriguingly, PCMD-1 was consistently more powerful to recruit SPD-5 and PLK-1 to the plasma membrane in the absence of an endogenous PCMD-1. In the *pcmd-1(t3421)* mutant background, SPD-5 and PLK-1 are not efficiently recruited to the centrosome and are expected to be more abundant in the cytoplasm. This finding suggests a ‘tug-of-war’ between the centrosomal and membrane-bound PCMD-1 pools for the recruitment of SPD-5 and PLK-1. Since PCMD-1 can also efficiently recruit SPD-5^4A^, a phospho-deficient version of SPD-5 for residues playing a key role in the expansion of the mitotic PCM scaffold, we speculate that the membrane-targeted SPD-5 represents the non-mitotic core.

In contrast to the PCM proteins, PCMD-1 was unable to relocate centriolar proteins SAS-4 and SAS-7 to the plasma membrane. This is consistent with the observations that the centriolar localization of both SAS-4 and SAS-7 is independent of PCMD-1 (this study and Erpf et al., 2019). Especially in the case of SAS-4, despite the strong protein-protein interaction in the yeast-two hybrid system, membrane-bound PCMD-1 is insufficient to overcome the ‘drag’ to the centriole probably because SAS-4 is stably incorporated in the centriole structure (Dammermann et al., 2008; Balestra et al., 2015).

In turn, the outer centriolar protein SAS-7 is required for PCMD-1 centrosomal recruitment in early embryos. However, in the yeast two-hybrid assay, we did not detect an interaction between PCMD-1 and SAS-7 or SPD-2. The interaction between SAS-7 and PCMD-1 is either indirect or an additional co-factor or modification is required for physical interaction in the yeast two-hybrid assay. The interaction between PCMD-1 and SAS-7 could be mediated through binding to SAS-4. However, since SAS-4 foci are still present in *sas-7(or452)* mutant embryos, where no PCMD-1 is found at the centrosome, we do not favor this possibility (Sugioka et al., 2017).

We conclude that a region in the C-terminal part of the protein spanning the last IDR is sufficient for the centrosomal localization of PCMD-1 but cannot sustain its function. The coiled-coil domain, which is located in the N-terminal part, is on its own insufficient for centrosomal targeting. However, the coiled-coil domain significantly contributes to the accumulation of PCMD-1 at the centrosome and is required to form an organized PCM. This could be a direct effect of the coiled-coil domain on SPD-5, as we have shown that PCMD-1 can interact with SPD-5, or an indirect effect due to the reduced PCMD-1 levels at the centrosome. In the yeast two-hybrid assay, we identified a strong self-interaction of PCMD-1, pointing to a tendency for oligomerization. Since this interaction is abolished in the absence of the coiled-coil domain, we hypothesize that the oligomerization of PCMD-1 through the coiled-coil domain plays an integral role in the PCM scaffold integrity.

We propose a model where SAS-7 recruits PCMD-1 to the centrioles. The C-terminal part of PCMD-1 is sufficient for its centrosomal recruitment. The coiled-coil domain, which is in the N-terminal part of the protein, enhances PCMD-1 accumulation through PCMD-1 oligomerization and thereby stabilizes the PCM scaffold. In turn, centrosomal PCMD-1 is sufficient to recruit the PCM scaffold protein SPD-5 and the mitotic kinase PLK-1 to the centrosome (Figure 6).

**Figure 6.**
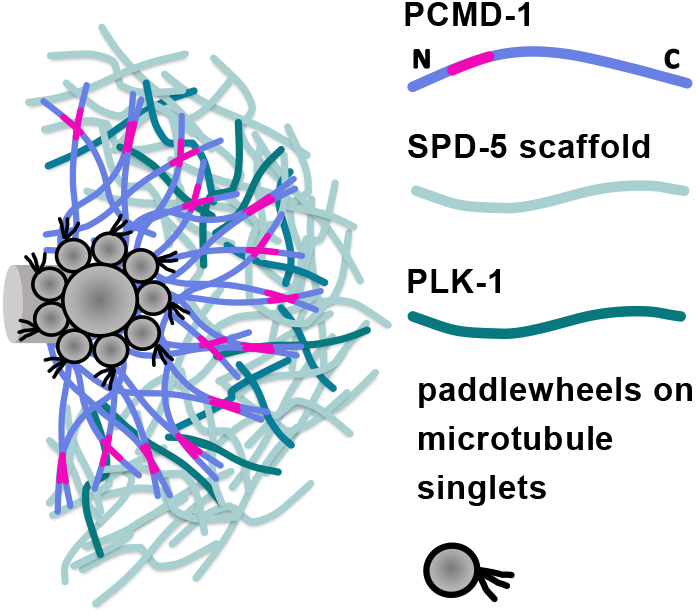
Model of how PCMD-1 bridges the centrioles and PCM. In the current model, PCMD-1 interacts with both centriolar and PCM proteins and thereby bridges the two centrosomal components. PCMD-1 is tethered to the centrioles through its C-terminal part, while the N-terminal part, including the coiled-coil domain, plays a role in PCMD-1 oligomerization and PCM formation.

In summary, we propose that PCMD-1 is a protein that directly anchors the PCM scaffold to the centrioles and thereby bridges the two centrosomal components.

## FIGURE LEGENDS

**Figure S1. SAS-7 recruits PCMD-1 to the centrioles in early embryos**

(A) Stills of time-lapse spinning disc confocal images of *gfp::pcmd-1* (n=6) and *gfp::pcmd-1;sas-7(or452)* (n=8) embryos during pronuclear migration. Centrosomes are shown enlarged for the GFP::PCMD-1 signal.

(B) Stills of time-lapse spinning disk confocal images of *gfp::pcmd-1* (n=7) and *gfp::pcmd-1;sas-7(or452)* (n=8) embryos during pronuclear meeting. Centrosomal areas were determined by DIC imaging and are shown enlarged for the GFP::PCMD-1 signal.

(C) Representative confocal images of fixed *gfp::pcmd-1* (n=3) and *gfp::pcmd-1;sas-7(or452)* (n=8) embryos (>6 nuclei) stained for DNA, GFP and SAS-4. Insets represent single channels of the centrioles.

In all panels, scale bars are 10 μm.

**Figure S2. PCMD-1 targeting of SPD-5 and PLK-1 to the plasma membrane in the presence of endogenous PCMD-1 is less efficient**

(A) Representative multicellular embryos of the ‘translocations assay’ using GFP::SPD-5 (n=19 no heat shock; n=19 heat shock), and PLK-1::sGFP (n=20 no heat shock; n=19 heat shock) in a wild-type background. Selected regions are enlarged and shown as merge and single channels. Note that plasma membrane-localized PLK-1::sGFP is faint. Scale bars are 10 μm.

(B) Quantification of (A); the percentage of embryos (%) with GFP signal at the membrane after heat shock in the wild-type background (n= total number of embryos analyzed).

**Figure S3. PCMD-1 is recruited before SPD-5 to sperm centrioles**

(A) Schematic representation of a marked mating experiment where *fog-2(n71)* females were mated with *gfp::pcmd-1* males (n=11). The image below represents a one-cell embryo taken by live-cell imaging. Centrosomal areas were determined by DIC imaging and are shown enlarged for the GFP::PCMD-1 signal.

(B) Schematic representation of a marked mating experiment where *fem-1* RNAi-treated *gfp::pcmd-1* females were mated with *fog-2(n71)* males (n=10). The image below represents a one-cell embryo taken by live-cell imaging. Centrosomal areas were determined by DIC imaging and are shown enlarged for the GFP::PCMD-1 signal.

(C) Images of fixed embryos in different stages of meiosis II, derived from the cross indicated in (B) and stained for DNA, GFP and SPD-5. Enlarged are sperm associated centrosomal signals merged and as single channels.

(D) Quantification of (C) percentage of embryos (%). n=number of analyzed embryos. Scale bars in all panels are 10 μm.

**Figure S4. The N-terminal region and the C1-region of PCMD-1 are not sufficient to target the protein to the centrosome**

(A) Embryonic survival (%) of *Pmai-2::gfp::pcmd-1* and *Pmai-2::gfp::pcmd-1(N)* in the *pcmd-1(t3421)* background at 15°C. Multiple Comparison with Kruskal Wallis test and post-hoc Dunn’s test adjusted with Holm correction, n=number of analyzed embryos.

(B) Images of embryos expressing *pcmd-1(t3421);Pmai-2::gfp::pcmd-1;Ppie-1::mCherry::h2b* (n=9), *pcmd-1(t3421);Pmai-2::gfp::pcmd-1(N);Ppie-1::mCherry::h2b* (n=7) and *pcmd-1(t3421);Pmai-2::gfp::pcmd-1(C1);Ppie-1::mCherry::h2b* (n=8) shortly after NEB. Centrosomes are shown enlarged for the GFP::PCMD-1 signal. n=number of analyzed embryos.

(C) Normalized cytoplasmic GFP signal intensities of embryos expressing *pcmd-1(t3421);Pmai-2::gfp::pcmd-1; Ppie-1::mCherry::h2b* (n=9, 1±0.005 a.u., mean±s.e.m.), *pcmd-1(t3421);Pmai-2::gfp::pcmd-1(N);Ppie-1::mCherry::h2b* (n=7, 1.42±0.04 a.u., mean±s.e.m.)*, pcmd-1(t3421);Pmai-2::gfp::pcmd-1(C1);Ppie-1::mCherry::h2b* (n=8, 1.14±0.0026 a.u., mean±s.e.m.) and *pcmd-1(t3421);Pmai-2::gfp::pcmd-1(C2);Ppie-1::mCherry::h2b* (n=9, 1.18±0.04 a.u., mean±s.e.m.) at NEB. n=number of analyzed embryos. Multiple comparison with ANOVA and post-hoc Tukey test.

(D) Localization of GFP::PCMD-1, GFP::PCMD-1(N), GFP::PCMD-1(C), GFP::PCMD-1(C1) and GFP::PCMD-1(C2) to the ciliary base in adult animals. n=5 animals for each condition.

In all panels error bars denote s.e.m.. P-values represent: **p<0.01, ****p<0.0001, ns= not significant. Scale bars are 10 μm.

**Movie 1. Time-lapse of the first cell cycle of a GFP::PCMD-1 expressing embryo** (related to figure 4C)

In the control embryos GFP::PCMD-1 localizes to the centrosome throughout the first cell cycle. Live-cell spinning disk microscopy. The scale bar is 10 μm.

**Movie 2. Time-lapse of the first cell cycle of a GFP::PCMD-1 expressing embryo lacking the predicted coiled-coil domain** (related to figure 4C)

In the *gfp::pcmd-1(ΔCC)* embryo GFP::PCMD-1(ΔCC) localizes to the centrosome with reduced levels. Live-cell spinning disk microscopy. The scale bar is 10 μm.

## METHODS

### *C. elegans* strains maintenance

Worms were maintained on NGM plates seeded with the OP50 *E. coli* strain under standard conditions at 15°C (Brenner, 1974). For experimental use worms were shifted to restrictive temperature of 25°C in L4 stage for 16-20h and progeny was imaged. *gfp::pcmd-1; sas-7(or452)/hT2* worms were allowed to lay eggs for 3h at 25°C. The laid eggs developed into adults at 25°C for 68h. Progeny of *gfp::pcmd-1;sas-7(or452)* worms were used for further analysis.

### Worm strain generation

Worms carrying single-copy transgene insertions were generated by the Universal MosSCI system, according to (Frøkjær-Jensen et al., 2008). Transgenes with the pCFJ350 backbone were injected into EG6699 or EG8081 and the progeny was selected using selection markers. Insertions were verified by PCR. Multiple independent insertion lines were screened for expression of the transgenes.

The *pcmd-1(syb1285 syb486[gfp::pcmd-1(ΔCC)])I* allele was generated by SunyBiotech by deleting 33 amino acids from E86 including F118, spanning the coiled-coil domain ranging from amino acid E86-N117, predicted by COILS program (see below). The deletion was verified by PCR and sequencing. The deletion was verified by PCR amplification using the oligos gcgctccgttgagaatctcgta and cacaaacgagcccgcacgga.

### Protein domain prediction and illustration

Intrinsically disordered regions were annotated based on information provided by the UniProt Consortium (UniProt Consortium, 2019). The coiled-coil domain was defined via the COILS program with a 28-residue window comparing both MTK and MTIDK matrices (weighted and unweighted) (Lupas et al., 1991; Lupas, 1996). The domain structures were illustrated using DOG2.0 (Ren et al., 2009).

### Yeast strains, media and transformation

Growth and genetic manipulation of the *Saccharomyces cerevisiae* strain EGY 48 (MATα, trp 1, his 3, ura 3, leu2::2/4/6 LECAop-LEU2) were performed using standard genetic techniques. The yeast strain was transformed with plasmids using lithium acetate (1M). The selection of the different plasmids was conducted with complete minimal medium lacking histidine/uracil/tryptophane/leucine. Yeast two-hybrid assays were performed using the Grow’N’Glow GFP Yeast Two-Hybrid System (Mobitech GmbH) according to the manufacturer’s protocol. Full-length cDNA of *C. elegans* SAS-7 and PCMD-1 were inserted into the bait vector pEG202 containing the DNA binding domain LexA. The cDNA of the different candidates SAS-7, SAS-4, PCMD-1, PCMD-1(ΔCC), SPD-2, SPD-5, PLK-1 were cloned into the prey vector pJG45 comprising the B42 transcription activation domain. The third plasmid transformed into the yeast was pGNG1-GFP containing the reporter gene *gfp*. pEG202-p53 with pJG45-LTA was used as a positive control, whereas pJG45 without an insert was used as a negative control. The presence of the plasmids in yeast was verified by plasmid extraction, followed by PCR amplification of the inserts and sequencing. The expression of the prey proteins was verified by immunoblotting against an HA-tag.

### SAS-4 feeding RNAi

A cDNA fragment comprising amino acids 2-455 was cloned into the L4440 vector and used to downregulate SAS-4 by feeding RNAi. Worms were incubated at 25°C for 28hs for partial and 30hs for full RNAi phenotypes.

### Translocation assay

L4 stage worms of different strains carrying the construct with the heat-shock promoter (TMD151, TMD157, TMD158, TMD159, TMD162, TMD165, TMD168, TMD167) were shifted to 25°C for 15h. Subsequently, worms were allowed to lay eggs for two hours at 25°C. The embryos were mounted on a 2 % agarose pad and heat-shocked at 30°C for 1h (Thermocycler Bio-Rad). After 2h recovery at 20°C, embryos were imaged at a SP5 Leica confocal microscope (see microscopy). Control embryos were incubated at 20°C without heat shock. Feeding RNAi was performed for 20h at 25°C by using I-2G08 for TMD168 and the pTMD118 feeding clone constructed against the reencoded region (Woodruff et al. 2015) for TMD151 and TMD162.

### Marked mating experiments

To mark the sperm centrioles in marked mating experiments *fog-2(q71),* females were mated with TMD119 males at 20°C and the progeny imaged by 4D-microscopy. For the converse experiment, TMD119 L4 worms were fed *fem-1(RNAi)* overnight. The hatched progeny was raised on *fem-1(RNAi)* to block sperm production. Feminized animals were mated with *fog-2(q71)* males at 20°C. Progeny of the crosses was either imaged with 4D-microscopy or used for indirect immunofluorescence.

### Indirect immunofluorescence

Indirect immunofluorescence was performed using a protocol by (Delattre et al., 2004). Hermaphrodite worms were cut in M9 buffer, covered with a coverslip and placed on ice blocks. After freeze-cracking, slides were fixed in methanol, followed by incubation with primary antibodies anti-SAS-4 (1:500, Santa Cruz Biotechnology), anti-SPD-5 (1:1000, a generous gift from B. Bowerman, Hamill et al., 2002) and anti-GFP (1:500, Roche) overnight at 4°C and with secondary antibodies Alexa488 (1:500, Invitrogen Molecular Probes), Alexa568 (1:500, Invitrogen Molecular Probes) and Hoechst 33258 (1:1000, Sigma) at room temperature for one hour.

### Microscopy

Embryos treated for the translocation assay and indirect immunofluorescence samples were imaged with a resolution of 1024×1024 pixels with HCX PL APO Lambda Blue 63x 1.4 oil objective and a step size of 0.7 μm at a SP5 Leica confocal microscope using the LAS software. For live-cell imaging young adult worms were either dissected in 6 μl M9 and mounted on 2% agar pads or dissected in Polybead^®^ Microspheres 20.00 μm (diluted 1:10 in M9). Live-cell imaging was performed on an inverted Nikon Eclipse Ti spinning disc confocal microscope using an Andor DU-888 X-11056 camera (1024 × 1024 pixels), a 100×1.45-NA Plan-Apochromat oil immersion objective and controlled by the NIS Elements 4.51 software. Z-stacks were taken every 30 s with a step size of 0.7 μm and with 2×2 binning. Embryos for marked mating experiments were imaged at the Zeiss Axio Imager.M2 equipped with epifluorescence and the Time to Live software from Caenotec. Differential interference contrast (DIC) 25 Z-stacks were taken throughout the volume of the embryo every 35 s, fluorescent scans were taken at required time points.

For structured illumination microscopy (SIM) of live *C. elegans* embryos, we used the ZEISS Elyra 7 system in the Lattice SIM mode equipped with a Plan Apochromat 63x/1.40 oil immersion objective. Images were acquired with two pco.edge sCMOS 4.2 cameras simultaneously, using the ZEISS DuoLink adapter. Acquired Z-stacks with a voxel size of 63×63×110 nm^3^ and FOV size of 1024×1024 pixels were processed using the SIM processing algorithm of ZEN Black 3.0 software. SIM processed Z-stacks have a voxel size of 31.5×31.5×110 nm^3^. Yeast colonies were acquired using a Leica Stereomicroscope M205 FA, controlled by the Leica Application Suite software (3.2.0.9652) and equipped with a 1x 2.11 NA Plan Apo lens and a Leica DigitalDFC340x FX camera.

### GFP intensity measurements

Centrosomal GFP intensities were measured on raw images at the time point of NEB by analyzing Z-stacks with ManualTrackMate in Fiji (Tinevez et al., 2017). The time point was either defined through the DNA condensation visualized by the mCherry::H2B marker (for TMD165, TMD166 and TMD177) or through corresponding DIC recordings (for PHX1285, TMD119). When there was no GFP signal recognizable, the centrosomal position was determined by DIC. A fixed radius of 0.788μm was applied to measure the GFP signal, background signal, and cytoplasmic background signal in 3D. Intensities were calculated for each centrosome: intensity = (C-B) – (CS-B). The total intensity of the background (B) was subtracted from the total intensity of the centrosome (C) and from the total intensity of the cytoplasmic signal (CS). The cytoplasmic signal without background was then subtracted from the centrosomal signal without background.

Statistical analysis was performed by using R Studio version 1.2.5003. Shapiro-Wilk test was used to test for normality. Levene’s test was performed to compare variances. Dependent on the normality, variance, and number of groups in the data sets, different comparisons tests were performed (see Figure legend). Mean values with the standard error of mean were plotted in Prism v6.

### Circularity measurements

Centrosomal circularities were evaluated in one-cell embryos ranging from NEB to metaphase that were immunostained with an antibody against SPD-5. The cell stage was defined by DNA condensation, visualized with Hoechst staining. Image analysis was performed in Fiji (Schindelin et al., 2012). Maximum Z-projections were created and the PCM shapes were converted into black/white outlines using the ‘Huang’ threshold. Statistical analysis was performed by using R Studio version 1.2.5003. Shapiro-Wilk test was used to test for normality. Levene’s test was performed to compare variances. Kruskal-Wallis test was used for the comparison of circularity values. Mean values with the standard error of mean were plotted with Prism v6.

### Statistical analysis for embryonic survival

L4 worms were singled and maintained at the corresponding temperature; laid eggs and hatched worms were counted. Statistical analysis was performed in R Studio version 1.2.5003. Shapiro-Wilk test was used to test for normality. Levene’s test was used to test for homogeneity of variances. Dependent on the normality, variance, and number of groups in the data sets, different comparison tests were performed (see Figure legend).

## ACKNOWLEDGMENTS

We thank the laboratories of C. Osman and N. Wagener for help with the yeast two-hybrid experiments; M. Antoniolli, N. Sharma and M. Plotnikova for generating tools; A. Bezler for critical comments on the manuscript; the imaging facility CALM, especially H. Harz and J. Ryan for help with imaging, the whole Zeiss-team and especially M. Gorelashvili for giving us the opportunity to image using the Lattice SIM, N. Lebedeva and C. Nöcker for excellent technical assistance, P. Gönczy for providing worm strains, B. Bowerman for the anti-SPD-5 antibody, the LSM graduate school. Some strains used in this study were provided by the Caenorhabditis Genetic Center (CGC), which is funded by the NIH Office of Research Infrastructure Programs (P40 OD010440). This work was supported by the German Research Council (DFG MI 1867/3).

## AUTHORS CONTRIBUTION

Experimental design, Methodology – LS, TMD; Resources, Investigation – LS, JM, AS, SÜ, EZu, TMD; Validation, Formal analysis, Supervision, Visualization, Writing – original draft preparation – LS, TMD; Writing – review and editing – LS, TMD, EZa; Conceptualization - TMD, EZa; Project administration, Funding acquisition -TMD.

## REFERENCES

Balestra, F.R., L. von Tobel, and P. Gönczy. 2015. Paternally contributed centrioles exhibit exceptional persistence in C. elegans embryos. Cell Res. 25:642–644. doi:10.1038/cr.2015.49.

Boxem, M., Z. Maliga, N. Klitgord, N. Li, I. Lemmens, M. Mana, L. de Lichtervelde, J.D. Mul, D. van de Peut, M. Devos, N. Simonis, M.A. Yildirim, M. Cokol, H.-L. Kao, A.-S. de Smet, H. Wang, A.-L. Schlaitz, T. Hao, S. Milstein, C. Fan, M. Tipsword, K. Drew, M. Galli, K. Rhrissorrakrai, D. Drechsel, D. Koller, F.P. Roth, L.M. Lakoucheva, A.K. Dunker, R. Bonneau, K.C. Gunsalus, D.E. Hill, F. Piano, J. Tavernier, S. van den Heuvel, A.A. Hyman, and M. Vidal. 2008. A Protein Domain-Based Interactome Network for C. elegans Early Embryogenesis. Cell. 134:534–545. doi:10.1016/j.cell.2008.07.009.

Brenner, S. 1974. The genetics of Caenorhabditis elegans. Genetics. 77:71–94.

Cabral, G., T. Laos, J. Dumont, and A. Dammermann. 2019. Differential Requirements for Centrioles in Mitotic Centrosome Growth and Maintenance. Developmental Cell. 1–34. doi:10.1016/j.devcel.2019.06.004.

Dammermann, A., P.S. Maddox, A. Desai, and K. Oegema. 2008. SAS-4 is recruited to a dynamic structure in newly forming centrioles that is stabilized by the gamma-tubulin-mediated addition of centriolar microtubules. The Journal of Cell Biology. 180:771–785. doi:10.1083/jcb.200709102.

Dammermann, A., T. Müller-Reichert, L. Pelletier, B. Habermann, A. Desai, and K. Oegema. 2004. Centriole Assembly Requires Both Centriolar and Pericentriolar Material Proteins. Developmental Cell. 7:815–829. doi:10.1016/j.devcel.2004.10.015.

Decker, M., S. Jaensch, A. Pozniakovsky, A. Zinke, K.F. O’Connell, W. Zachariae, E. Myers, and A.A. Hyman. 2011. Limiting Amounts of Centrosome Material Set Centrosome Size in C.elegans Embryos. Current Biology. 21:1259–1267. doi:10.1016/j.cub.2011.06.002.

Delattre, M., C. Canard, and P. Gönczy. 2006. Sequential Protein Recruitment in C. elegans Centriole Formation. Current Biology. 16:1844–1849. doi:10.1016/j.cub.2006.07.059.

Delattre, M., S. Leidel, K. Wani, K. Baumer, J. Bamat, H. Schnabel, R. Feichtinger, R. Schnabel, and P. Gönczy. 2004. Centriolar SAS-5 is required for centrosome duplication in C. elegans. Nature Cell Biology. 6:656–664. doi:10.1038/ncb1146.

Erpf, A.C., L. Stenzel, N. Memar, M. Antoniolli, M. Osepashvili, R. Schnabel, B. Conradt, and T. Mikeladze-Dvali. 2019. PCMD-1 Organizes Centrosome Matrix Assembly in C.elegans. Current Biology. 29:1324–1336.e6. doi:10.1016/j.cub.2019.03.029.

Fields, S., and O. Song. 1989. A novel genetic system to detect protein-protein interactions. Nature. 340:245–246. doi:10.1038/340245a0.

Frøkjær-Jensen, C., M.W. Davis, C.E. Hopkins, B.J. Newman, J.M. Thummel, S.-P. Olesen, M. Grunnet, and E.M. Jorgensen. 2008. Single-copy insertion of transgenes in Caenorhabditis elegans. Nat Genet. 40:1375–1383. doi:10.1038/ng.248.

Hamill, D.R., A.F. Severson, J.C. Carter, and B. Bowerman. 2002. Centrosome maturation and mitotic spindle assembly in C. elegans require SPD-5, a protein with multiple coiled-coil domains. Developmental Cell. 3:673–684. doi: 10.1016/s1534-5807(02)00327-1.

Hilbert, M., M.C. Erat, V. Hachet, P. Guichard, I.D. Blank, I. Flückiger, L. Slater, E.D. Lowe, G.N. Hatzopoulos, M.O. Steinmetz, P. Gönczy, and I. Vakonakis. 2013. Caenorhabditis elegans centriolar protein SAS-6 forms a spiral that is consistent with imparting a ninefold symmetry. Proc. Natl. Acad. Sci. U.S.A. 110:11373–11378. doi:10.1073/pnas.1302721110.

Kemp, C.A., K.R. Kopish, P. Zipperlen, J. Ahringer, and K.F. O’Connell. 2004. Centrosome maturation and duplication in C. elegans require the coiled-coil protein SPD-2. Developmental Cell. 6:511–523. doi: 10.1016/s1534-5807(04)00066-8.

Kirkham, M., T. Müller-Reichert, K. Oegema, S. Grill, and A.A. Hyman. 2003. SAS-4 Is a C. elegans Centriolar Protein that Controls Centrosome Size. Cell. 112:575–587. doi:10.1016/s0092-8674(03)00117-x.

Kitagawa, D., I. Vakonakis, N. Olieric, M. Hilbert, D. Keller, V. Olieric, M. Bortfeld, M.C. Erat, I. Flückiger, P. Gönczy, and M.O. Steinmetz. 2011. Structural Basis of the 9-Fold Symmetry of Centrioles. Cell. 144:364–375. doi:10.1016/j.cell.2011.01.008.

Leidel, S., and P. Gönczy. 2003. SAS-4 is essential for centrosome duplication in C. elegans and is recruited to daughter centrioles once per cell cycle. Developmental Cell. 4:431–439. doi:10.1016/s1534-5807(03)00062-5.

Leidel, S., M. Delattre, L. Cerutti, K. Baumer, and P. Gönczy. 2005. SAS-6 defines a protein family required for centrosome duplication in C. elegans and in human cells. Nature Cell Biology. 7:115–125. doi:10.1038/ncb1220.

Lettman, M.M., Y.L. Wong, V. Viscardi, S. Niessen, S.-H. Chen, A.K. Shiau, H. Zhou, A. Desai, and K. Oegema. 2013. Direct Binding of SAS-6 to ZYG-1 Recruits SAS-6 to the Mother Centriole for Cartwheel Assembly. Developmental Cell. 25:284–298. doi:10.1016/j.devcel.2013.03.011.

Letunic, I., and P. Bork. 2018. 20 years of the SMART protein domain annotation resource. Nucleic Acids Research. 46:D493–D496. doi:10.1093/nar/gkx922.

Li, S., C.M. Armstrong, N. Bertin, H. Ge, S. Milstein, M. Boxem, P.-O. Vidalain, J.-D.J. Han, A. Chesneau, T. Hao, D.S. Goldberg, N. Li, M. Martinez, J.-F. Rual, P. Lamesch, L. Xu, M. Tewari, S.L. Wong, L.V. Zhang, G.F. Berriz, L. Jacotot, P. Vaglio, J. Reboul, T. Hirozane-Kishikawa, Q. Li, H.W. Gabel, A. Elewa, B. Baumgartner, D.J. Rose, H. Yu, S. Bosak, R. Sequerra, A. Fraser, S.E. Mango, W.M. Saxton, S. Strome, S. van den Heuvel, F. Piano, J. Vandenhaute, C. Sardet, M. Gerstein, L. Doucette-Stamm, K.C. Gunsalus, J.W. Harper, M.E. Cusick, F.P. Roth, D.E. Hill, and M. Vidal. 2004. A map of the interactome network of the metazoan C. elegans. Science. 303:540–543. doi:10.1126/science.1091403.

Lupas, A., Van Dyke, M., and Stock, J. 1991. Predicting Coiled Coils from Protein Sequences Science 252:1162–1164. DOI: 10.1126/science.252.5009.1162

Lupas, A. 1996 .Prediction and Analysis of Coiled-Coil Structures Meth. Enzymology 266:513–525. DOI: 10.1016/s0076-6879(96)66032-7

McNally, K.L.P., A.S. Fabritius, M.L. Ellefson, J.R. Flynn, J.A. Milan, and F.J. McNally. 2012. Kinesin-1 Prevents Capture of the Oocyte Meiotic Spindle by the Sperm Aster. Developmental Cell. 22:788–798. doi:10.1016/j.devcel.2012.01.010.

Mittasch, M., V.M. Tran, M.U. Rios, A.W. Fritsch, S.J. Enos, B. Ferreira Gomes, A. Bond, M. Kreysing, and J.B. Woodruff. 2020. Regulated changes in material properties underlie centrosome disassembly during mitotic exit. The Journal of Cell Biology. 219:647–23. doi:10.1083/jcb.201912036.

O’Connell, K.F., C. Caron, K.R. Kopish, D.D. Hurd, K.J. Kemphues, Y. Li, and J.G. White. 2001. The C. elegans zyg-1 gene encodes a regulator of centrosome duplication with distinct maternal and paternal roles in the embryo. Cell. 105:547–558. doi: 10.1016/s0092-8674(01)00338-5.

Pelletier, L., E. O’Toole, A. Schwager, A.A. Hyman, and T. Müller-Reichert. 2006. Centriole assembly in Caenorhabditis elegans. Nature. 444:619–623. doi:10.1038/nature05318.

Pelletier, L., N. Özlü, E. Hannak, C. Cowan, B. Habermann, M. Ruer, T. Müller-Reichert, and A.A. Hyman. 2004. The Caenorhabditis elegans Centrosomal Protein SPD-2 Is Required for both Pericentriolar Material Recruitment and Centriole Duplication. Current Biology. 14:863–873. doi:10.1016/j.cub.2004.04.012.

Qiao, R., G. Cabral, M.M. Lettman, A. Dammermann, and G. Dong. 2012. SAS-6 coiled-coil structure and interaction with SAS-5 suggest a regulatory mechanism in C. elegans centriole assembly. EMBO J. 1–14. doi:10.1038/emboj.2012.280.

Ren, J., L. Wen, X. Gao, C. Jin, Y. Xue, and X. Yao. 2009. DOG 1.0: illustrator of protein domain structures. Cell Res. 19:271–273. doi:10.1038/cr.2009.6.

Rogala, K.B., N.J. Dynes, G.N. Hatzopoulos, J. Yan, S.K. Pong, C.V. Robinson, C.M. Deane, P. Gönczy, and I. Vakonakis. 2015. The Caenorhabditis elegans protein SAS-5 forms large oligomeric assemblies critical for centriole formation. eLife. 4:e07410. doi:10.7554/eLife.07410.

Schindelin, J., I. Arganda-Carreras, E. Frise, V. Kaynig, M. Longair, T. Pietzsch, S. Preibisch, C. Rueden, S. Saalfeld, B. Schmid, J.-Y. Tinevez, D.J. White, V. Hartenstein, K. Eliceiri, P. Tomancak, and A. Cardona. 2012. Fiji: an open-source platform for biological-image analysis. Nat Meth. 9:676–682. doi:10.1038/nmeth.2019.

Schultz, J., R.R. Copley, T. Doerks, C.P. Ponting, and P. Bork. 2000. SMART: a web-based tool for the study of genetically mobile domains. Nucleic Acids Research. 28:231–234. doi:10.1093/nar/28.1.231.

Sugioka, K., D.R. Hamill, J.B. Lowry, M.E. McNeely, M. Enrick, A.C. Richter, L.E. Kiebler, J.R. Priess, and B. Bowerman. 2017. Centriolar SAS-7 acts upstream of SPD-2 to regulate centriole assembly and pericentriolar material formation. eLife. 6:1–25. doi:10.7554/elife.20353.

Tinevez, J.-Y., N. Perry, J. Schindelin, G.M. Hoopes, G.D. Reynolds, E. Laplantine, S.Y. Bednarek, S.L. Shorte, and K.W. Eliceiri. 2017. TrackMate: An open and extensible platform for single-particle tracking. Methods. 115:80–90. doi:10.1016/j.ymeth.2016.09.016.

UniProt Consortium. 2019. UniProt: a worldwide hub of protein knowledge. Nucleic Acids Research. 47:D506–D515. doi:10.1093/nar/gky1049.

Varadarajan, R., and N.M. Rusan. 2018. Bridging centrioles and PCM in proper space and time. Essays in Biochemistry. 62:793–801. doi:10.1042/EBC20180036.

Woodruff, J.B., B.F. Gomes, P.O. Widlund, J. Mahamid, A. Honigmann, and A.A. Hyman. 2017. The Centrosome Is a Selective Condensate that Nucleates Microtubules by Concentrating Tubulin. Cell. 169:1066–1071.e10. doi:10.1016/j.cell.2017.05.028.

Woodruff, J.B., O. Wueseke, V. Viscardi, J. Mahamid, S.D. Ochoa, J. Bunkenborg, P.O. Widlund, A. Pozniakovsky, E. Zanin, S. Bahmanyar, A. Zinke, S.H. Hong, M. Decker, W. Baumeister, J.S. Andersen, K. Oegema, and A.A. Hyman. 2015. Regulated assembly of a supramolecular centrosome scaffold in vitro. Science. 348:808–812. doi:10.1126/science.aaa3923.

